# Quantification of DNA samples by Ethidium Bromide Spot Technique

**DOI:** 10.1101/289108

**Authors:** Jessica Zhao, Julian Zhao, Antoine Cummins, Tiffany Gonzalez, Grace Axler-DiPerte, Devin Camenares

## Abstract

Accurate and quick determination of DNA concentration is critical for the assembly of synthetic constructs, as well as a multitude of other experiments. We sought to optimize an under-utilized and inexpensive approach for determining DNA concentration: a spotting technique that uses the intercalating dye Ethidium Bromide. This technique does not require specialized equipment such as a spectrophotometer, but instead relies on visualization of dye-DNA complex fluorescence when excited by UV light. We modelled and tested a range of parameters for dye concentration and spot size, finding that 15uL spots with 1.0ug/mL Ethidium Bromide produced the most reliable standard curve. More importantly, we hope that our approach can help other labs optimize this protocol for their own experimental setup. Adoption of this technique may help enable development of iGEM teams in resource limited environments and laboratories which do not or cannot employ a satisfactory method for determining DNA concentration.

**Financial Disclosure:** We obtained funding support from Canon Solutions America, and indirect support for this work through the CUNY Research Scholars Program. However, the funders had no role in study design, data collection and analysis, decision to publish, or preparation of the manuscript.

**Competing Interests:** The authors have declared that no competing interests exist.

**Ethics Statement:** N/A

**Data Availability:** Yes – all data are fully available without restriction. The data can be found in the associated supplementary materials.

**Article Tags:** DNA, Ethidium Bromide, EtBr, Concentration, Method, UV, Gibson Assembly

**Header Statement:** **This work was assessed during the iGEM/PLOS Realtime Peer Review Jamboree on 23rd February 2018 and has been revised in response to the reviewers.** Responses to Reviewers, and supplementary information (including Fig. S1) are available as separate files.

## Introduction

Successful completion of many synthetic biology experiments, and most (if not all) iGEM projects, involve some amount of DNA assembly. This may be conventional restriction enzyme cloning, such as 3A assembly, or more powerful strategies such as Gibson assembly^1^. Regardless of the method used, accurate quantification of the concentration of DNA samples used is often critical to success. For example, improper ratios of DNA fragments in Gibson assembly and similar techniques can lead to a dramatically reduced efficiency^2^.

A variety of equipment exists to provide accurate and reliable measurements of DNA concentration in a sample. Devices such as the NanoDrop require a very small amount of a DNA sample to provide information on the amount and purity contained therein. However, the NanoDrop and even older, less sophisticated spectrophotometers can be prohibitively expensive. For example, the retail price of a NanoDrop can exceed ten thousand dollars; even a used UV-VIS spectrophotometer, which may not be as reliable, can still cost in excess of five hundred dollars. While these expenses and equipment requirements may not be an obstacle for iGEM teams at institutions where extensive research is a high priority, this can represent a significant challenge for teams with limited resources, such as a community college team.

We tried to address the problem of DNA quantitation during the course of our project, primarily to aid our assembly step. One alternative to the use of a spectrophotometer is to run a small amount of a DNA sample on an agarose gel which is subsequently stained with Ethidium Bromide (EtBr), an intercalating agent that is relatively cheap^3,4^. EtBr emits red/orange light in response to excitation by UV light, and emission is greatly increased through binding DNA, probably because unbound dye fluorescence is quenched by water^5^. The intensity of the fluorescence from a EtBr – DNA complex is proportional to the length and amount of DNA present in a band on an agarose gel.

While agarose gel electrophoresis is routinely used to assess the size of DNA fragments in a sample, we thought it was too laborious and time consuming for routine quantification of the amount in a sample. Instead, we decided to employ a variation on this theme, and simply spot the DNA and EtBr together on a solid surface. This spot technique has been described before, but we could not find a published study that systematically explored and optimized the protocol to our satisfaction^6^.

In this report, we describe our efforts to standardize a protocol for measuring DNA concentration by the EtBr spot technique. It is our hope that future iGEM teams can follow our protocol as well as our approach to optimize it. This may further enable the development of teams in community colleges, community labs, or other resource deprived settings that do not have access to spectrophotometers.

## Methods

### Modelling EtBr-DNA Fluorescence

A mass action reaction was used to model binding of EtBr to DNA. Using a dissociation constant^7^ (K_d_) of 15 s^−1^, the equation was rearranged to solve a particular amount of EtBr-DNA complex for given starting concentration of EtBr and DNA (Fig. S1). This algebraic rearrangement was expedited by using the Wolfram Alpha webserver (https://www.wolframalpha.com/) using the query “solve for c k = ((a- c)(b-c))/c” in which C is the complex, A is the dye, B is the DNA, and K is the dissociation constant. Fluorescence of the complex was assumed to be 20-fold brighter than the dye alone^5^; the detection range for EtBr was determined experimentally and used to estimate the detectable range of EtBr-DNA complex.

### Ethidium Bromide Spotting Method

5-15uL of diluted Ethidium Bromide (Bio-Rad) was pipetted onto a disposable plastic petri dish. 1uL of water or diluted lambda DNA (New England Biolabs) was then pipetted into the droplet/spot containing the dye and mixed. The dish was then inverted over a UV transilluminator. A cover shielded ambient light and a photograph was taken using an orange light filter and an android smartphone (Motorola Droid 2).

### Image Processing

Images of the spots were processed using ImageJ^8^. Briefly, spots were selected using a freehand or oval tool, either to analyze the entire spot (Whole Area) or the region within the spot with the brightest uniform intensity (Highest Fluorescence). Pixel intensity was measured using the analysis tool, with or without a de-speckling correction applied first (Noise reduction). These measurements were either taken as a ratio to a background signal and/or normalized to the maximum value for a set, and averaged across multiple experiments.

## Results

We wished to determine what ratio and amounts of EtBr and DNA would generate the most useful standard curve for a given range of concentrations. This will depend not only on the range for which a linear relationship is desired, but also on the detection limits for EtBr in our experimental setup. Towards that end, we tested different concentrations of EtBr alone, from 100 ng/uL to 5 pg/uL. From these results, we determined that our detection range for EtBr is approximately 1ng/uL to 10ng/uL; anything below this range produces no measurable fluorescence, while concentration differences above this range cannot be easily distinguished.

Given information on fluorescence of EtBr alone and our range of detection, we were able to model the detection of EtBr-DNA complexes, which are approximately 20-fold brighter than the free dye molecule. These results (Fig 1) suggest that for DNA concentrations in the range of 10 – 100 ng/uL (amounts typical for DNA assembly), EtBr concentrations of 0.5 ng/uL to 5ng/uL work best. Below this level, we predict that there is not enough EtBr to detect, and a significant fraction of DNA remains unbound by EtBr. On the contrary, too much EtBr should produce a significant amount of background signal and likewise diminish the capacity to quantitate DNA samples.

**Fig 1:**
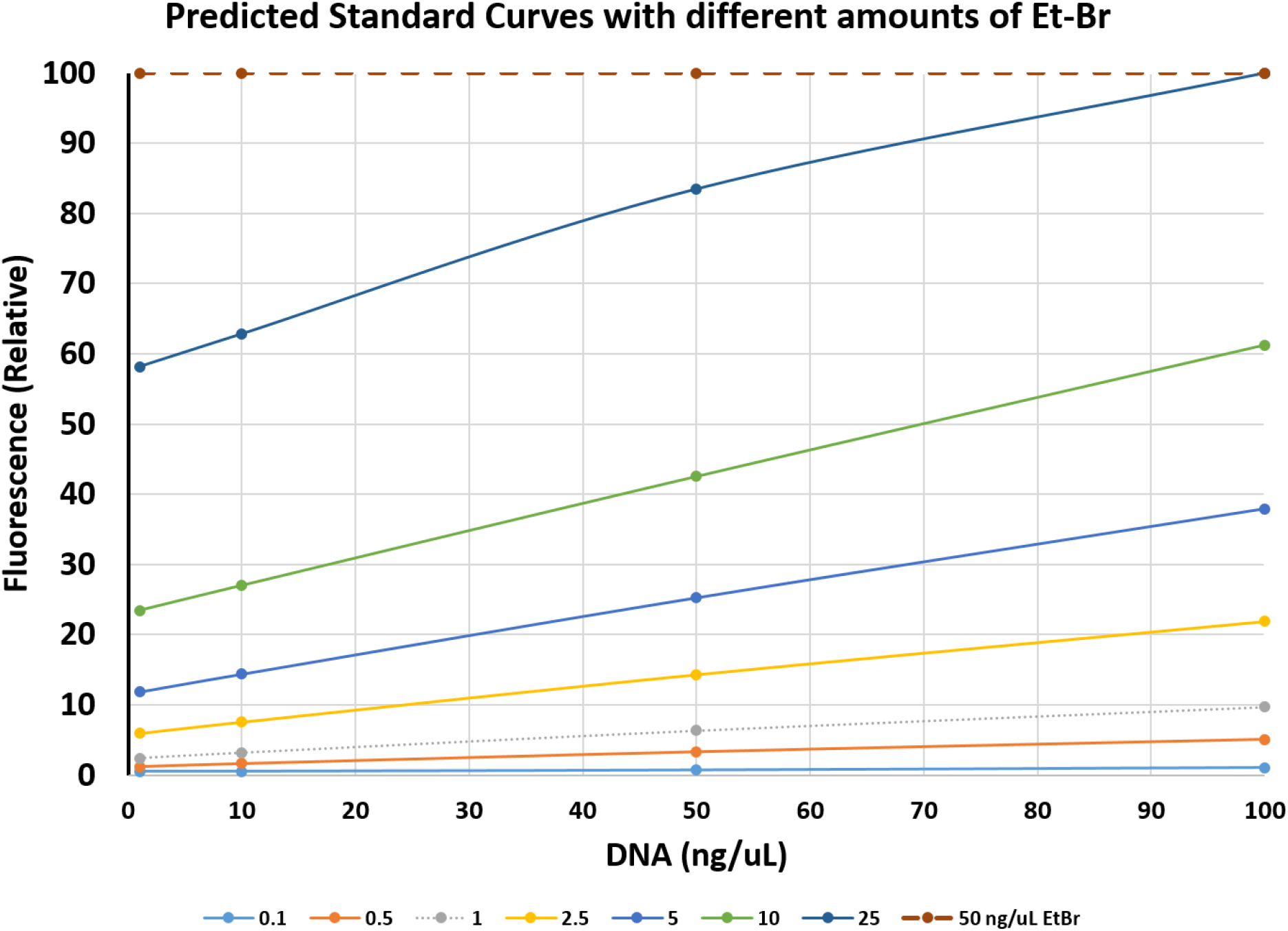
Predicted detection of fluorescence for given DNA and Ethidium Bromide concentrations. Binding of Ethidium Bromide to DNA was modelled for different concentrations of each molecule. Based on measurement of the dye alone, the predicted values were normalized to the maximum detectable fluorescence. Different color lines indicate fluorescence of different concentrations of dye across a range of different concentrations of DNA (x-axis).

We proceeded to test this model by spotting different concentrations of lambda DNA together with EtBr (Fig 2). Since the concentration of DNA in these samples are known, we can create standard curves for each amount of EtBr or total volume used. The ideal standard curve will present a gradual and smooth increase in pixel intensity versus DNA concentration – conditions that create such a curve are therefore useful in determining the concentration of an unknown sample. We used different approaches for processing the image, achieving the best curve through eliminating noise and limiting measurements to a region of each spot with uniform intensity (Fig 3). This approach was used to compare curves using 0.5ng/uL, 1.0ng/uL, or 2.5ng/uL of EtBr with a consistent drop size (Fig 4), in which the intermediate amount of fluorescence gave the best curve. This concentration of EtBr seemed to work better the larger the droplet size, with 15uL working the best and 5uL performing the worst (Fig 5).

**Fig 2:**
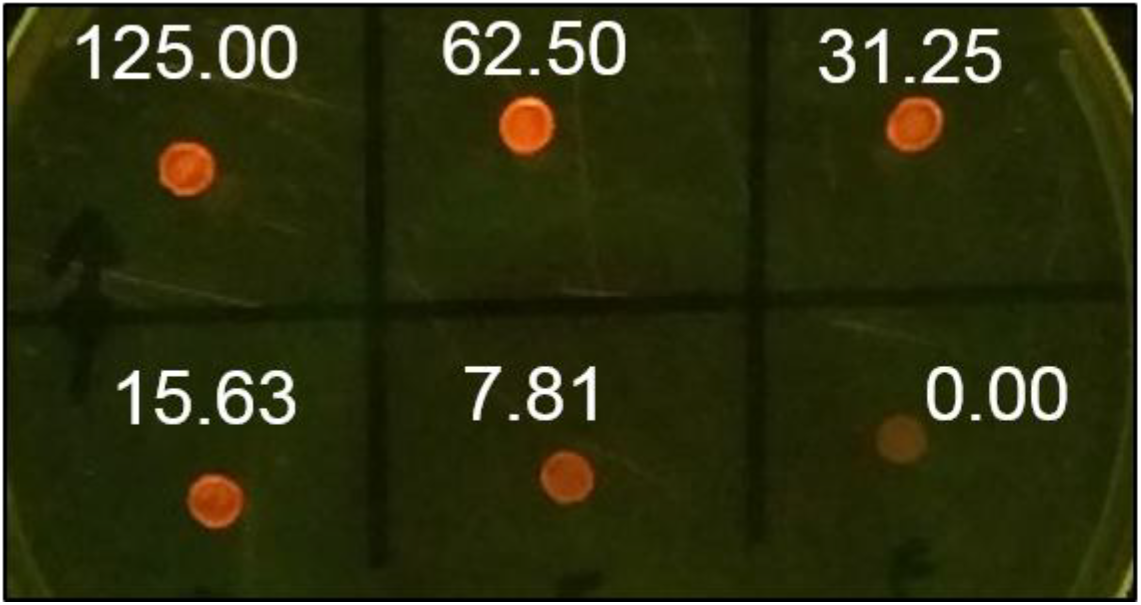
Fluorescence of Ethidium Bromide – DNA spots. Spots containing 10uL of 0.5ng/uL Ethidium Bromide and 1uL of a lambda DNA standard were illuminated and photographed as described in the methods. The concentration of DNA, in ng/uL, is shown above each spot in white text. This is a representative image of several experiments and concentrations.

**Fig 3:**
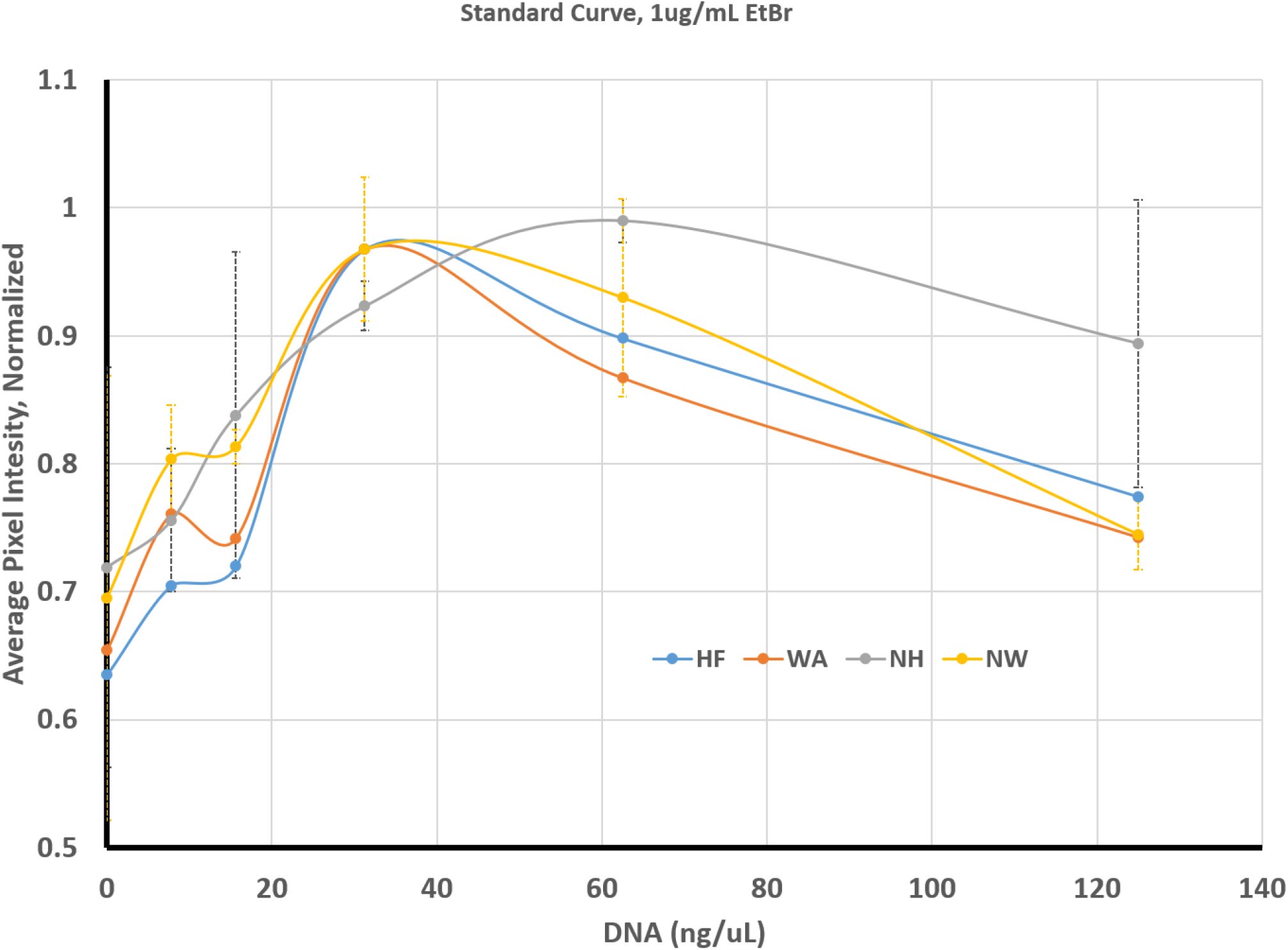
Standard Curve for DNA concentration versus Pixel Intensity with 1.0ug/mL. Droplets were prepared and measured as indicated in the methods. The measurements were taken for each droplet as either the area of highest uniform fluorescence (HF, Blue LIne), the same area with noise reduction (NH, Gray Line), the whole area of the droplet (WA, Orange Line), or this whole area with noise reduction (NW, Yellow Line). The data was normalized for each replicate against the highest value obtained and plotted as a fraction of this maximum value. Error bars represent the standard deviation from averaging three independent replicates, with the color of the error lines corresponding to the colors of the lines. Only error bars for the noise reduced highest fluorescence and whole area.

**Fig 4:**
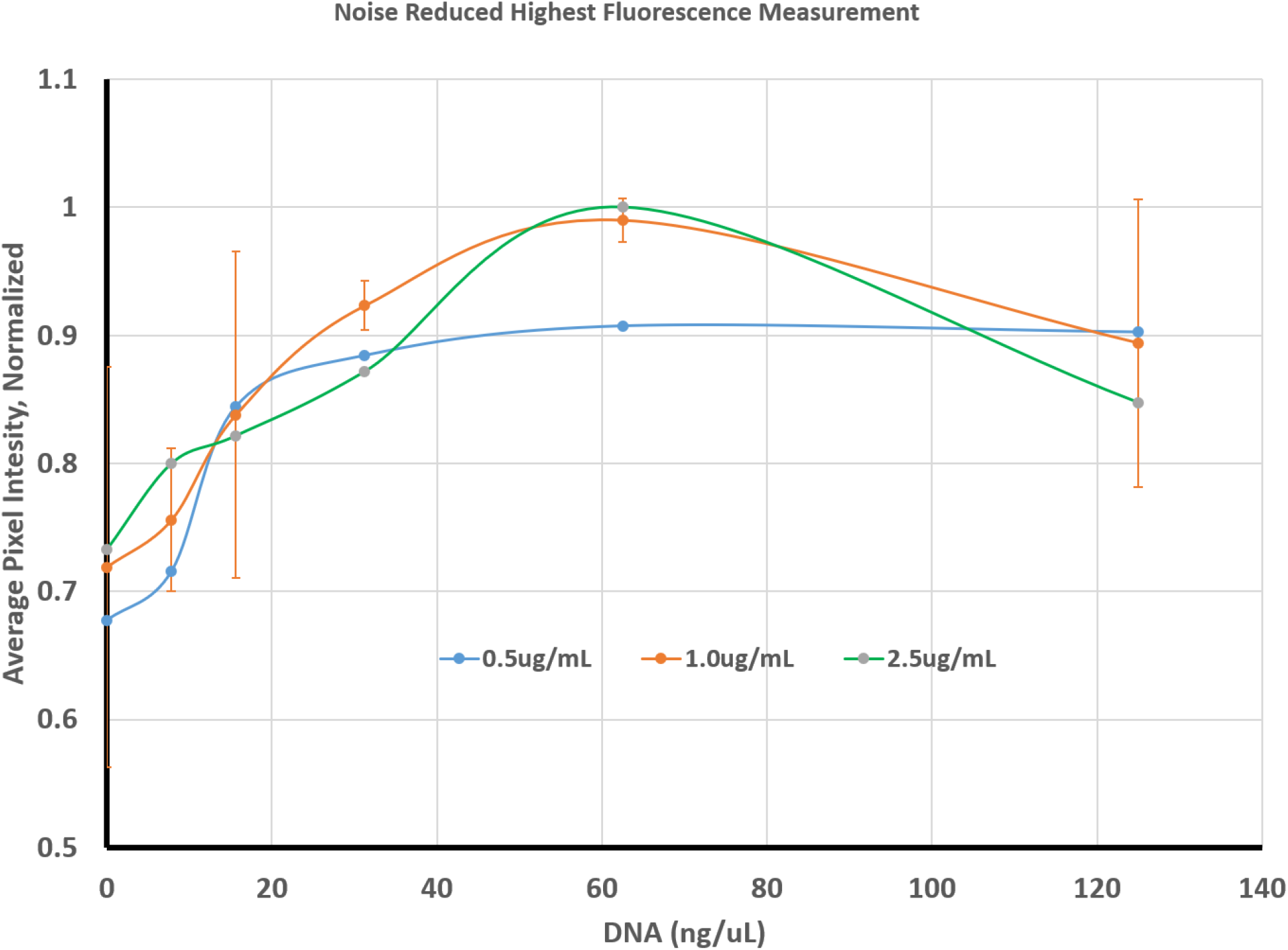
Comparison of different amounts of Ethidium Bromide. Droplets were prepared and measured as indicated in the methods; images were processed by eliminating noise and analyzing only the area of highest uniform fluorescence within the droplet (same as NH in Fig 3). This was done using three different concentrations of Ethidium Bromide: 0.5ug/mL (Blue Line), 1.0ug/mL (Orange Line), or 2.5ug/mL (Green Line). The scale and error bars (for 1.0ug/mL only) are the same as described in Fig 3.

**Fig 5:**
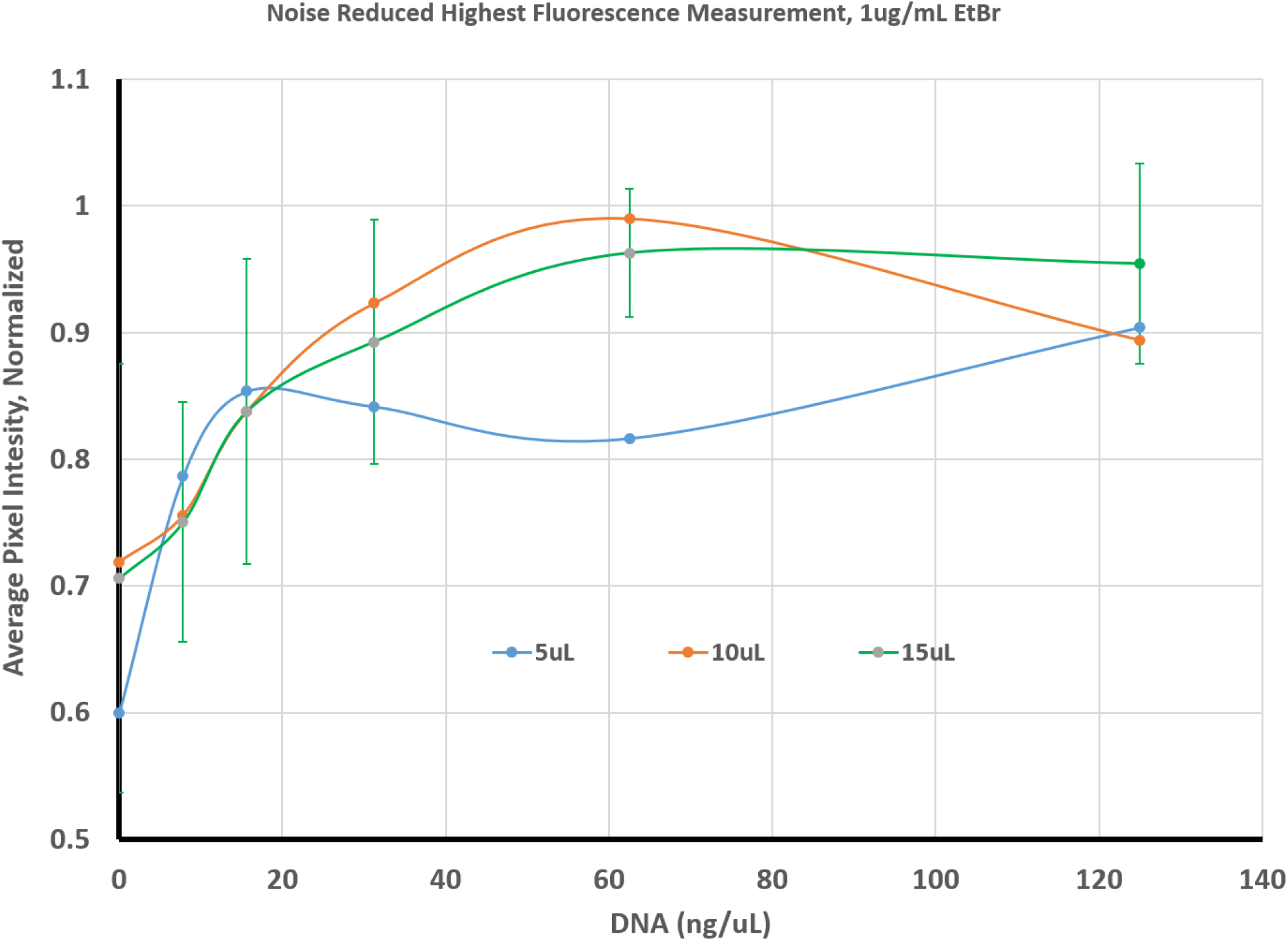
Comparison of Droplet Size. Droplets were prepared and measured as indicated in the methods; each droplet contained 1.0ug/mL of Ethidium Bromide and images were processed by eliminating noise and analyzing only the area of highest uniform fluorescence within the droplet (same as NH in Fig 3). Droplet size was varied between 5ul (Blue Line), 10uL (Orange Line), and 15uL (Green Line). The scale and error bars (for 15uL only) are the same as described in Fig 3.

## Discussion

In this report, we provide an approach for optimizing an inexpensive and easy DNA quantification protocol. This protocol or technique involves spotting EtBr onto a solid surface, mixing with a DNA sample, and observing fluorescence of the complex through illumination with UV light. We both modelled and tested how different amounts of DNA, different amounts of EtBr, and different size droplets affect the measurements and subsequent calculation of DNA concentration in a sample.

We found that in our experimental setup, the most reliable curves were ones that used a 15uL droplet with 1.0ug/mL of EtBr. Interestingly, our model would have predicted better results using a higher concentration, but in practice the best curve is arguably the one with the same conditions as previously described^6^. However, there was a great deal of variety between replicates, possibly due to human error. For example, a failure to properly mix the dye and DNA could lead to fluorescent intensity that is not uniform across the spot. Thus, it is advisable to measure a known DNA standard alongside every unknown sample that needs to be quantified. This is precisely the approach we used to aid our DNA assembly reactions, which were successful. Alternatively, different dilutions (1:2, 1:4) of an unknown DNA sample can be measured simultaneously to increase the accuracy of the quantification.

It is our hope that our approach will be useful for future iGEM teams from community colleges, or other settings that lack conventional equipment for measuring DNA concentration. They will be able to follow the model we set forth here and determine the optimal conditions for using the EtBr spot technique with their own experimental setup. Future attempts to optimize this protocol may include testing different types of cameras and filters. It should also be possible to create a more sophisticated optical model for account for the way in which the light will interact with and travel through different size spots. However, our preliminary results with varying spot size suggest that this will not be the most critical factor for generating useful data on DNA concentrations.

In future studies, we may explore alternatives to EtBr, since this intercalating dye may have toxic or carcinogenic properties. Alternative DNA-binding dyes such as SYBR Green (Life Technologies Co.) or GelRed (Biotium Inc.) are thought to be safer than EtBr yet still retain sufficient specificity and affinity for use in determining DNA concentration. The basic principles and protocol explored in this study should be easily adapted for any indicator dye that can be assessed visually.

Accurate measurement in synthetic biology is important not only for ensuring efficient DNA assembly, but also for determining if a design is functioning correctly. Wherever inexpensive and accessible alternatives to conventional approaches are possible, a wider range of aspiring synthetic biologists will be empowered to design, test and build. Likewise, we have also tried to adapt the spot imaging approach for measuring fluorescence of bacteria on an agar plate. Hopefully our approach to DNA quantification will find utility in other labs, and inspire other techniques that remove obstacles for those interested in participating in iGEM.

