## Supplementary Materials for "Quantification of DNA samples by Ethidium Bromide Spot Technique"

**Supplementary Data and Figures**

The images and measurements used in this paper are available as an excel file with embedded images, provided as supplementary data (igem_imgJ_replicates.xlsx). The measurements, processed data and calculations used for the graphs presented in this study are avaislable as another supplementary file (Average_data.xlsx). The equations in Fig S1 (next page) were generated using the following LaTeX format:

**A)** K_{d} = \frac{[E][D]}{[ED]}

**B)** K_{d} = \frac{([E_{0}]-[ED])([D_{0}]-[ED])}{[ED]}

**C)** [ED] = \frac{1}{2} \cdot (\sqrt{[E]^{2} - 2[E]([D]-K_{d})+([D]+K_{d})^{2}} + [E]+ [D] + K_{d})

These equations were employed in another excel file to predict the fluorescence that would be observed for varying amounts of Ethidium Bromide and DNA (EtBr_calculations.xlsx). This file also includes the measurements for EtBr fluorescence alone, with an embedded image of the data from which these measurements were taken.

**A**

$$K_d = \frac{[E][D]}{[ED]}$$

**B**

$$K_d = \frac{([E_0] - [ED])([D_0] - [ED])}{[ED]}$$

**C**

$$[ED] = \frac{1}{2} \cdot (\sqrt{[E]^2 - 2[E]([D] - K_d) + ([D] + K_d)^2} + [E] + [D] + K_d)$$

**Fig. S1**: **Mass-action equations model Ethidium Bromide – DNA binding. A)** The dissociation constant relates the concentration of Ethidium Bromide – DNA complex ([ED]) to remaining free Ethidium Bromide ([E]) and DNA ([D]). In these equations, DNA concentration is modelled not based upon the actual size of DNA used, but rather on available sets of 2 base pairs (4 bases) upon which the dye can bind. **B)** A substituted equation in which final or equilibrium concentrations of either the dye or DNA is described as a function of the initial concentration (i.e. $[E_0]$ as the initial concentration of the dye) minus the equilibrium levels of the complex. **C)** A rearranged version of the mass-action equation in which the amount of complex ([ED]) is solvable given the starting parameters of initial dye and DNA (here just [E] and [D]).
