## Supplementary Materials for "Quantification of DNA samples by Ethidium Bromide Spot Technique"

Author Contributions:

Conceptualization: JuZ, DC

Methodology: JeZ, JuZ

Investigation: JeZ, JuZ, AnC

Formal Analysis: JeZ, JuZ, TG

Writing – Original Draft: DC, JeZ

Writing – Review & Editing: GAD, DC, JuZ, JeZ

Abstract

Accurate and quick determination of DNA concentration is critical for the assembly of synthetic constructs, as well as a multitude of other experiments. We sought to optimize an under-utilized and inexpensive approach for determining DNA concentration: a spotting technique that uses the intercalating dye Ethidium Bromide. This technique does not require specialized equipment such as a spectrophotometer, but instead relies on visualization of dye-DNA complex fluorescence when excited by UV light. We modelled and tested a range of parameters for dye concentration and spot size, finding that 15uL spots with 1.0ug/mL Ethidium Bromide produced the most reliable standard curve. More importantly, we hope that our approach can help other labs optimize this protocol for their own experimental setup. Adoption of this technique for determining DNA concentration may help enable development of iGEM teams in resource limited environments.

**Comments**

**lgallagher:** "More importantly, we hope that our approach can help other labs optimize this protocol for their own experimental setup. Adoption of this technique for determining DNA concentration may help enable development of iGEM teams in resource limited environments." Explaining how this research could be used in future studies is a great point to have in your Abstract as it clearly explains the benefits of your research and who a key readership might be.

Financial Disclosure

We obtained funding support from Canon Solutions America, and indirect support for this work through the CUNY Research Scholars Program. However, the funders had no role in study design, data collection and analysis, decision to publish, or preparation of the manuscript.

Competing Interests

The authors have declared that no competing interests exist.

Ethics Statement

N/A

Data Availability

Yes – all data are fully available without restriction. The data can be found in the associated supplementary materials.

Article Tags

DNA, Ethidium Bromide, EtBr, Concentration, Method, UV, Gibson Assembly

*** Full Submission ***

Introduction:

Successful completion of many synthetic biology experiments, and most (if not all) iGEM projects, involve some amount of DNA assembly. This may be conventional restriction enzyme cloning, such as 3A assembly, or more powerful strategies such as Gibson assembly1. Regardless of the method used, accurate quantification of the concentration of DNA samples used is often critical to success. For example, improper ratios of DNA fragments in Gibson assembly and similar techniques can lead to a dramatically reduced efficiency2.

A variety of equipment exists to provide accurate and reliable measurements of DNA concentration in a sample. Devices such as the NanoDrop require a very small amount of a DNA sample to provide information on the amount and purity contained therein. However, the NanoDrop and even older, less sophisticated spectrophotometers can be prohibitively expensive. For example, the retail price of a NanoDrop can exceed ten thousand dollars; even a used UV-VIS spectrophotometer, which may not be as reliable, can still cost in excess of five hundred dollars. While these expenses and equipment requirements may not be an obstacle for iGEM teams at research-1 university, this can represent a significant challenge for teams with limited resources, such as a community college team.

**Comments**

**ChrisFerguson:** 'research-1 university' is a US-centric term - perhaps avoid these sorts of terms especially as you're hoping to appeal globally to folks in resource-poor settings.

We tried to address the problem of DNA quantitation during the course of our project, primarily to aid our assembly step. One alternative to the use of a spectrophotometer is to run a small amount of a DNA sample on an agarose gel which is subsequently stained with Ethidium Bromide (EtBr), an intercalating agent that is relatively cheap3,4. EtBr emits red/orange light in response to excitation by UV light, and emission is greatly increased through binding DNA, probably because unbound dye fluorescence is quenched by water5. The intensity of the fluorescence from a EtBr – DNA complex is proportional to the length and amount of DNA present in a band on an agarose gel.

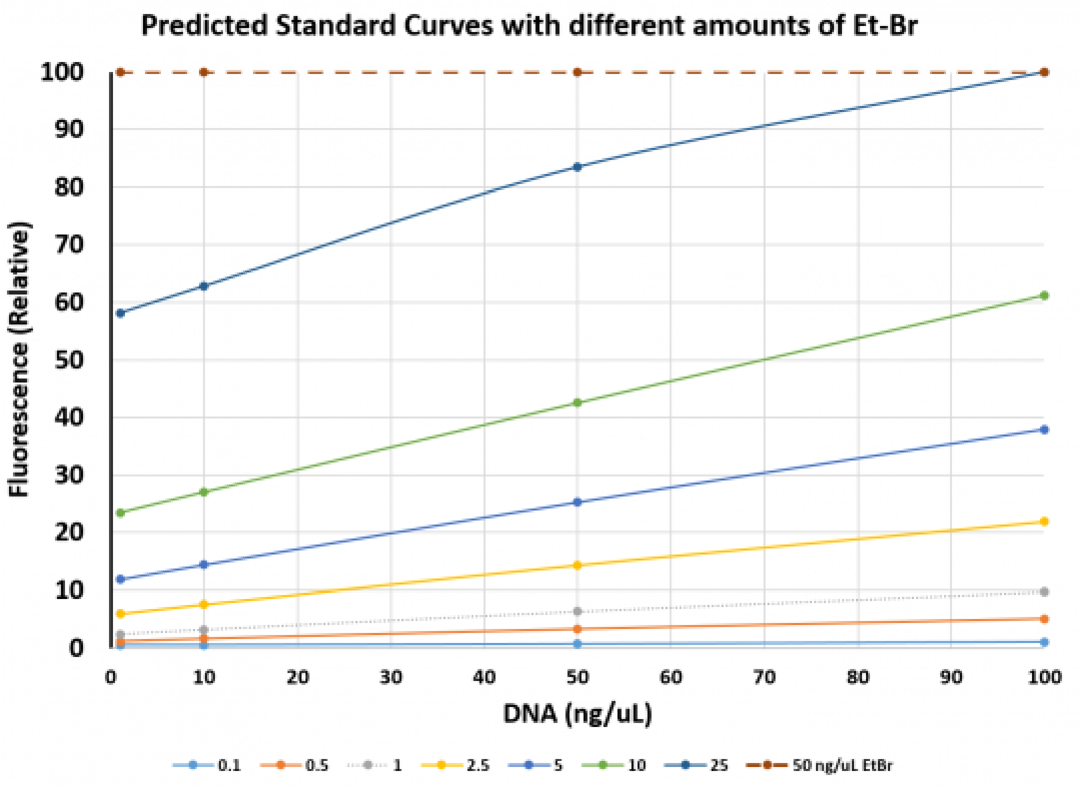


Fig 1: Predicted detection of fluorescence for given DNA and Ethidium Bromide concentrations. Binding of Ethidium Bromide to DNA was modelled for different concentrations of each molecule. Based on measurement of the dye alone, the predicted values were normalized to the maximum detectable fluorescence. Different color lines indicate fluorescence of different concentrations of dye across a range of different concentrations of DNA (x-axis).

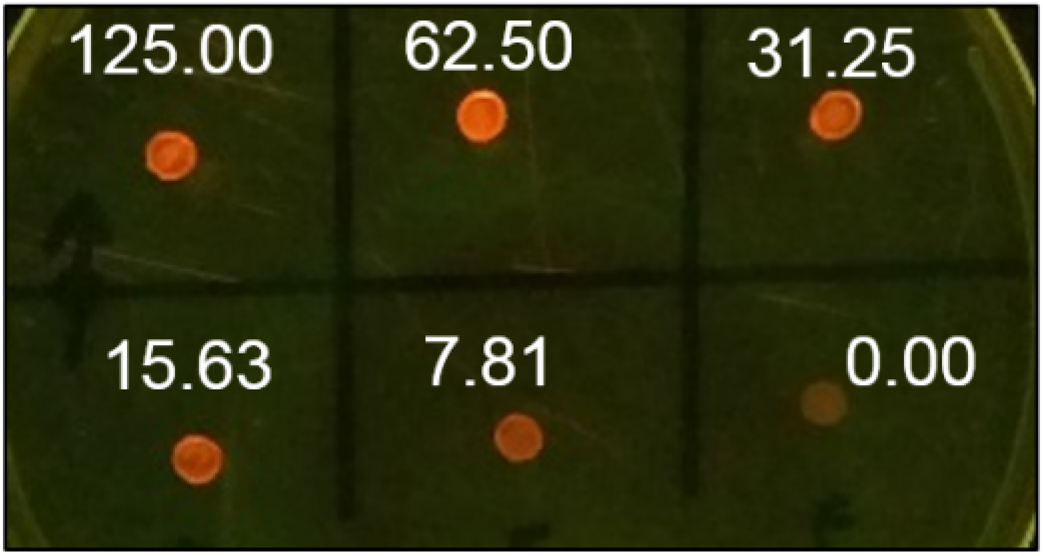


Fig 2: Fluorescence of Ethidium Bromide – DNA spots. Spots containing 10uL of 0.5ng/uL Ethidium Bromide and 1uL of a lambda DNA standard were illuminated and photographed as described in the methods. The concentration of DNA, in ng/uL, is shown above each spot in white text. This is a representative image of several experiments and concentrations.

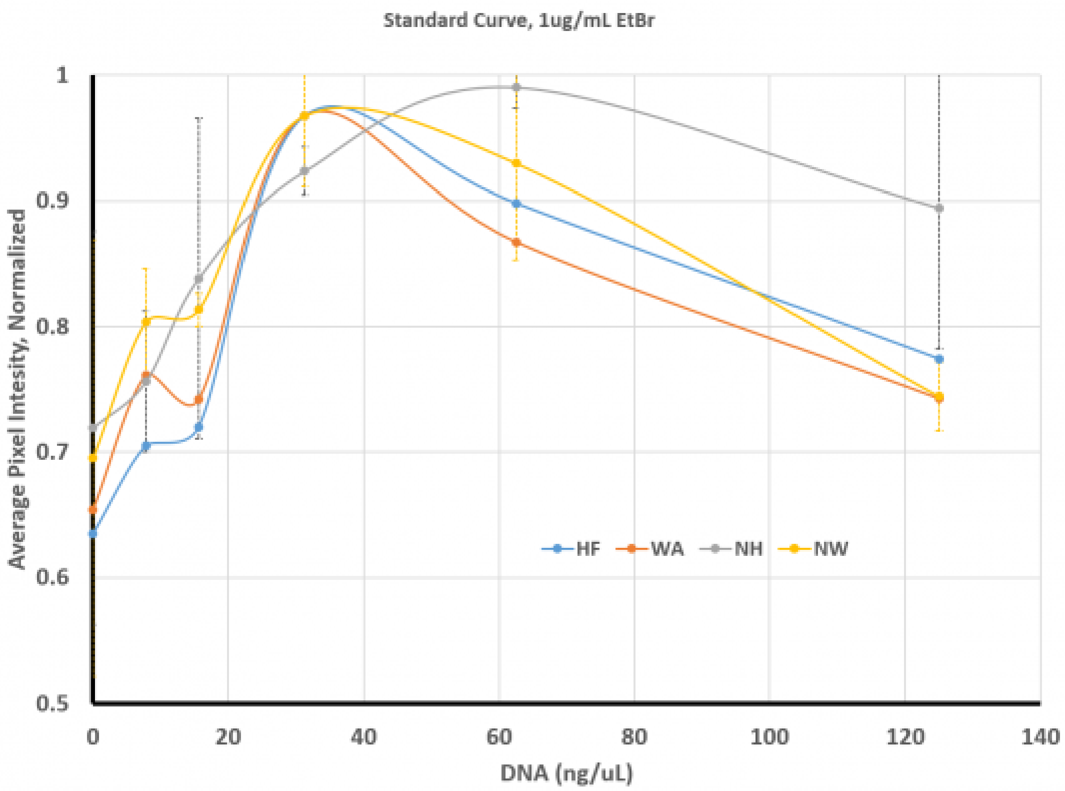


Fig 3: Standard Curve for DNA concentration versus Pixel Intensity with 1.0ug/mL. Droplets were prepared and measured as indicated in the methods. The measurements were taken for each droplet as either the area of highest uniform fluorescence (HF, Blue LIne), the same area with noise reduction (NH, Gray Line), the whole area of the droplet (WA, Orange Line), or this whole area with noise reduction (NW, Yellow Line). The data was normalized for each replicate against the highest value obtained and plotted as a fraction of this maximum value. Error bars represent the standard deviation from averaging three independent replicates, with the color of the error lines corresponding to the colors of the lines. Only error bars for the noise reduced highest fluorescence and whole area.

**Comments**

**lgallagher:** Hi City University of New York, Thanks for your submission. It looks as though the error bars for figures 3, 4 and 5 have been cut off at the top, maybe these could be reformated to ensure these pull onto the graphs?

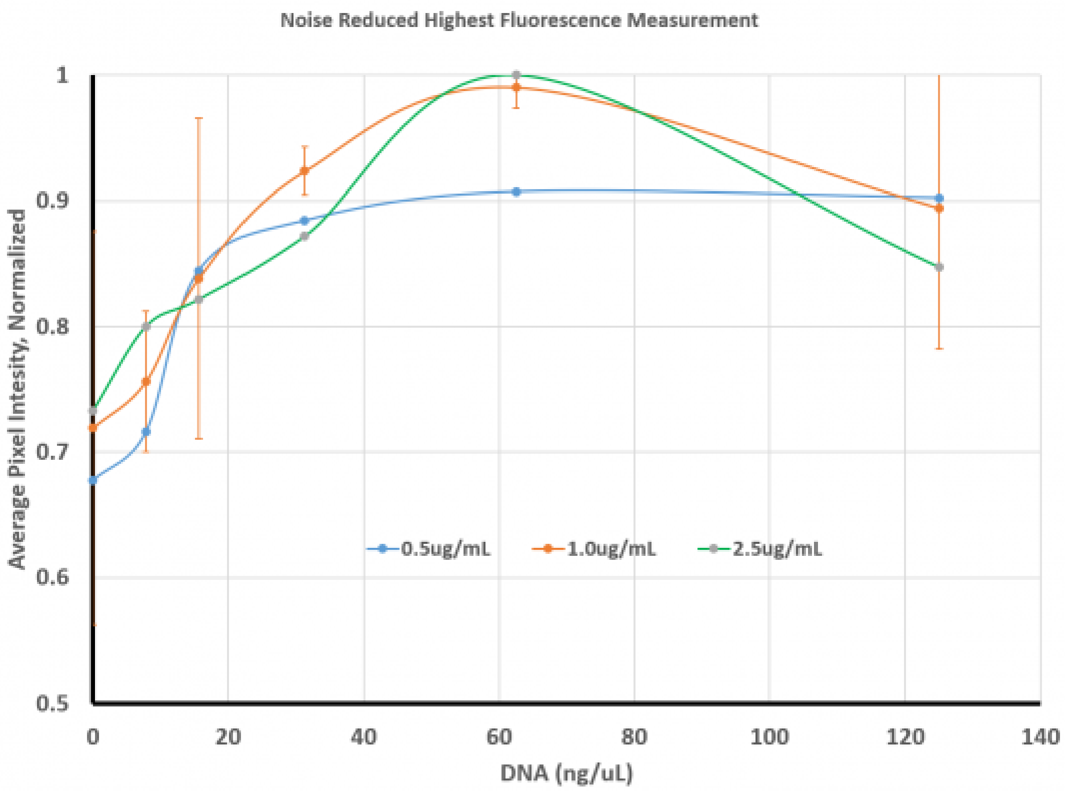


Fig 4: Comparison of different amounts of Ethidium Bromide. Droplets were prepared and measured as indicated in the methods; images were processed by eliminating noise and analyzing only the area of highest uniform fluorescence within the droplet (same as NH in Fig 3). This was done using three different concentrations of Ethidium Bromide: 0.5ug/mL (Blue Line), 1.0ug/mL (Orange Line), or 2.5ug/mL (Green Line). The scale and error bars (for 1.0ug/mL only) are the same as described in Fig 3.

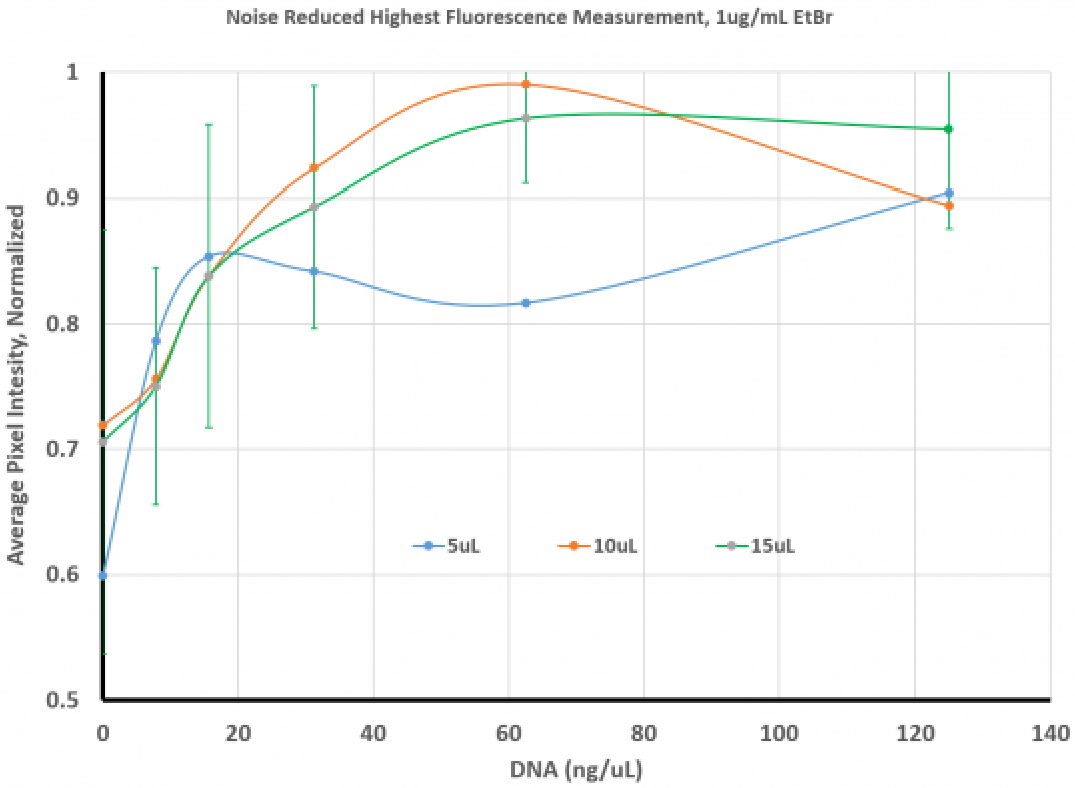


Fig 5: Comparison of Droplet Size. Droplets were prepared and measured as indicated in the methods; each droplet contained 1.0ug/mL of Ethidium Bromide and images were processed by eliminating noise and analyzing only the area of highest uniform fluorescence within the droplet (same as NH in Fig 3). Droplet size was varied between 5ul

(Blue Line), 10uL (Orange Line), and 15uL (Green Line). The scale and error bars (for 15uL only) are the same as described in Fig 3.

**A**

$$K_d = \frac{[E][D]}{[ED]}$$

**B**

$$K_d = \frac{([E_0] - [ED])([D_0] - [ED])}{[ED]}$$

**C**

$$[ED] = \frac{1}{2} \cdot (\sqrt{[E]^2 - 2[E]([D] - K_d) + ([D] + K_d)^2} + [E] + [D] + K_d)$$

Fig. S1: Mass-action equations model Ethidium Bromide – DNA binding. A)The dissociation constant relates the concentration of Ethidium Bromide – DNA complex ([ED]) to remaining free Ethidium Bromide ([E]) and DNA ([D]). In these equations, DNA concentration is modelled not based upon the actual size of DNA used, but rather on available sets of 2 base pairs (4 bases) upon which the dye can bind. B) A substituted equation in which final or equilibrium concentrations of either the dye or DNA is described as a function of the initial concentration (i.e. [E0] as the initial concentration of the dye) minus the equilibrium levels of the complex. C) A rearranged version of the mass-action equation in which the amount of complex ([ED]) is solvable given the starting parameters of initial dye and DNA (here just [E] and [D]).

Supplementary Data and Information

The images and measurements used in this paper are available as an excel file with embedded images, provided as supplementary data (igem_imgJ_replicates.xlsx). The measurements, processed data and calculations used for the graphs presented in this study are avaislable as another supplementary file (Average_data.xlsx). The equations in Fig S1 were generated using the following LaTeX format:

**Comments**

**lgallagher:** Hi City University of New York, Please could you upload your Supporting Information file or provide a link to this?
**Djcamenares:** Sorry for the inconvenience - must not have been uploaded correctly upon submission. Here are the files (zipped) attached, also accessible via DropBox: https://www.dropbox.com/s/pb7nzxub87cqa7v/PLOS_One_iGEM_KBCC_Supplementary.zip?dl=0

A) K_{d} = \frac{[E][D]}{[ED]}

B) K_{d} = \frac{([E_{0}]-[ED])([D_{0}]-[ED])}{[ED]}

C) [ED] = \frac{1}{2} \cdot (\sqrt{[E]^{2} - 2[E]([D]-K_{d})+([D]+K_{d})^{2}} + [E]+ [D] + K_{d})

These equations were employed in another excel file to predict the fluorescence that would be observed for varying amounts of Ethidium Bromide and DNA (EtBr_calculations.xlsx). This file also includes the measurements for EtBr fluorescence alone, with an embedded image of the data from which these measurements were taken.

**Comments**

**lgallagher:** Hi City University of New York, Considering the potential risks associated with Ethidium Bromide (it is a mutagenic substance), have you considered any possible alternative dyes, and if so, which ones and how would you apply them?

**Djcamenares:** Hi Larch - Great suggestion. It is something we considered but did not explore or mention. There are alternatives, like GelRed or GelGreen, which can be substituted, probably in a similar way. Of course, this would involve using different dissociation constants, and starting from scratch with simulating and then obtaining the results. We will incorporate mention of them in a revised draft.

**ChrisFerguson:** Hi Devin and colleagues, Useful contribution! Have you considered uploading your optimised detection protocol on protocols.io and then including a link in the methods. It's a great way to ensure the protocol is cited and you get the credit - it also encourages users to record improvements which would get layered onto the original protocol. If you haven't already - check out the possibilities here: https://www.protocols.io/

> **Djcamenares:** Great suggestion - definitely something we will do (with this protocol) and probably will use for other methods hereafter. Thanks
