## Supplementary Materials for "Quantification of DNA samples by Ethidium Bromide Spot Technique"

**Response to Reviewers**

The comment **lgallagher** by regarding the abstract does not necessarily suggest a revision – the impact of the use of the technique in future studies is stated, as the reviewer highlights. However, in order to clarify our meaning, we reworded the final sentence of the abstract: Adoption of this technique may help enable development of iGEM teams in resource limited environments *and laboratories which do not or cannot employ a satisfactory method for determining DNA concentration.*

We followed the recommendation by **ChrisFerguson** and changed the phrase “research-1 university” to “institutions where extensive research is a high priority”, which is an appropriate definition for research-1 university and in line with the actual classification given by the Carnegie Foundation for the Advancement of Teaching.

We also followed the recommendation by **lgallagher** regarding Figure 3, 4, and 5; we fixed the axis of these graphs accordingly.

We also included a paragraph in the discussion to mention the use of alternatives to Ethidium Bromide, as suggested by **lgallagher**

We did not upload our protocol to https://www.protocols.io/ as suggested by **ChrisFerguson**, since the parameters we used may have depended on our specific equipment and setup. Although we may upload such a protocol in the future, we recommend that other laboratories followed our approach and optimize the parameters for the protocols for their own equipment.
